# Pleiotropic Variability Score: A Genome Interpretation Metric to Quantify Phenomic Associations of Genomic Variants

**DOI:** 10.1101/2021.07.18.452819

**Authors:** Khader Shameer, Benjamin S. Glicksberg, Marcus A. Badgeley, Kipp W. Johnson, Joel T. Dudley

## Abstract

A more complete understanding of phenomic space is critical for elucidating genome-phenome relationships and for assessing disease risk from genome sequencing. To incorporate knowledge of how related a variant’s associations are, we developed a new genome interpretation metric called Pleiotropic Variability Score (PVS). PVS uses semantic reasoning to score the relatedness of a genetic variant’s associated phenotypes based on those phenotypes’ relationships in the human phenotype ontology (HPO) and disease ontology (DO). We tested 78 unique semantic similarity methods and integrated six robust metrics to define the pleiotropy score of SNPs. We computed PVS for 12,541 SNPs which were mapped to 382 HPO and 317 DO unique phenotype terms in a genotype-phenotype catalog (10,021 SNPs mapped to DO phenotypes and 8,569 SNPs mapped to HPO phenotypes). We validated the utility of PVS by computing pleiotropy using an electronic health record linked genomic database (Bio*ME*, n=11,210). Further we demonstrate the application of PVS in personalized medicine using “personalized pleiotropy score” reports for individuals with genomic data that could potentially aid in variant interpretation. We further developed a software framework to incorporate PVS into VCF files and to consolidate pleiotropy assessment as part of genome interpretation pipelines. As the genome-phenome catalogs are growing, PVS will be a useful metric to assess genetic variation to find SNPs with highly pleiotropic effects. Additionally, variants with varying degree of pleiotropy can be prioritized for explorative studies to understand specific roles of SNPs and pleiotropic hubs in mediating novel phenotypes and drug development.

## INTRODUCTION

Phenomics is the systematic study of the physical and biochemical traits of organisms due to variation in genetic, metabolic, environmental or other perturbations^1, 2^. Phenomics is an emerging component of multiscale biology which seeks to improve the understanding of the pathophysiology of human disease to advance translational science^3^ and aid drug discovery^4^. Evaluation of biological or clinical phenotypes (phenome) will help in ascertaining the genotype-phenotype relationships mediating complex diseases and genetic disorders^5^. The translational bioinformatics community provides tools and methods to capture, mine, and analyze phenomics data^6–8^. With the recent growth in human genome-phenome databases, pleiotropy can be compiled from by systematic integration of phenotypes associated with a genetic loci^9, 10^. Phenomic inference tools (PheGenI^11^, PhenoGen^12^), ontologies^13^ and databases (PhenomicDB^14^) for capturing and integrating biological and clinical phenomic data are expanding. Recent efforts to unify human phenotype knowledge into ontologies have fuelled quantitative estimates of human phenotypes to predict disease modalities^15, 16^. Electronic medical record (EMR) linked genome-wide and phenome-wide association studies (GWAS and PheWAS) expanded our understanding of how a single genotype could influence multiple phenotypes. Variant-specific, trait-specific, or systematic PheWAS has uncovered novel genotype-phenotype associations that further show the network architecture of human diseases at the genome-phenome level. For example, genetic variants affecting platelet traits were associated with various conditions including inflammatory spondylopathies, hemorrhoids, acute bronchospasm, and myocardial infarction^17–19^.

Multiple studies have shown that rare, orphan, complex, or genetic diseases have shared genomic and phenomic components^20–23^. “Pleiotropy”^24^ is the phenomenon of a genetic variant, gene or specific genomic region within an linkage disequilibrium block influencing multiple phenotypic traits and plays an important role in the evolution of traits and complex phenotypes^25–29^. Studying the network properties of human genes in the context of pleiotropic effects reveals the role of one gene in different disease pathogeneses^30, 31^. For example, 8q24^32^ is associated with prostate cancer, breast cancer, ovarian cancer, colorectal cancer and bladder cancer whereas 9p21^33, 34^ is associated multiple cardiovascular diseases. While these regions affect distinct clinical manifestations - they are mediated by an underlying common mechanism of pathophysiology (e.g. cancers and disease of cardiovascular systems) and specific genomic regions showing hypermutation (kataegis) trends^35^.

Pleiotropy on a SNP level is often more apparent. For example, rs3184504, a trypsin to arginine variant on an adapter protein SH2B3, is associated with more than 12 phenotypes including hypothyroidism, blood pressure, type 1 diabetes, coronary heart disease, arthritis, myocardial infarction, peripheral arterial disease and several medical traits like eosinophil counts^19, 36^. On the other end of the pleiotropy spectrum, an intronic SNP within *BTBD9*, rs9357271, is only associated with restless leg syndrome^37^. Here the SNPs have several phenomic associations, but a quantification of pleiotropy is not possible without the understanding of individual diseases and their etiology by manual abstraction of disease classes using manual curation. In a typical genome-interpretation workflow, the variants with multiple phenotypes get prioritized irrespective of how similar or correlated the phenotypes are. Quantifying phenomic similarity of a given variant can better prioritize variants with pleiotropic effects than counting the number of phenotype associations.

Genomic variant interpretation methods help to understand the contributions of SNPs or structural variants to the clinical manifestation of a complex diseases ^38, 39^. Existing metrics for variant interpretations are based on gene or protein sequence information, sequence conservation, evolutionary divergence, and perturbation impact of variant on the protein structure^38^. These measures provide possible links between the clinical phenotype and variant data generated from whole genome sequencing (WGS) or whole exome sequencing (WES) ^40, 41^. Current methods do not estimate the similarity or diversity of the phenotype set associated with a given variant. Furthermore, current guidelines for genome interpretation or software frameworks do not consider phenomic associations of a genetic variant for prioritizing variants^39, 42, 43^. Despite the growth in translational biology and large-scale PheWAS studies^18, 44–47^, phenomics based metrics are sparse in variant interpretation^39, 42^.

In this report, we introduce a new metric called Pleiotropic Variability Score (PVS) to quantify the diversity of phenotypes associated with genetic variants. The method is developed using annotations compiled from a database of genotypes and phenotypes and two biomedical ontologies. A wide range of ontology based similarity assessment measures have been developed for information extraction from ontologies and structured data in the fields of biocuration^48^, natural language processing (NLP)^49^, artificial intelligence^50^, and Semantic Web mining^51^. Such semantic similarity scores were used as the algorithmic bases for analyzing massive biological and clinical datasets including functional similarity of gene ontology (GO) terms, comparative phenomics of plant species, gene prioritization, and disease network reconstruction^52–55^.

We have validated PVS as a metric using an EMR-linked genomics database (Bio*ME* biobank). We packaged the PVS system into a software framework (PVSAnnotate) to introduce PVS into VCF files as part of genome-annotation pipelines and to generate reports for genomic interpretations. Collectively our method, software, and the associated database for quantifying pleiotropy of genetic variants could provide a quantitative phenomics score as a component of genome interpretation pipelines.

## METHODS

We designed a scoring method to quantify pleiotropy of phenotypes associated with a variant. We used GWASdb v2 to define genotype-phenotype relationships and two ontologies to capture relationships between phenotypes. We developed the scoring method using a systematic survey of various semantic similarity measures and identify a combination of metrics that can effectively compute pleiotropy. Briefly, pleiotropy can be defined as the inverse of the semantic similarity of two biological entities (for example phenotypes) in a mathematical expression. This simple estimation gets complex due to the availability of multiple semantic similarity methods, and often a gene or an SNP may have more than two functions associated with GWAS, PheWAS, or function test experimental assays. Here we describe the ontologies, semantic similarity scores, and estimation used to develop a new method for quantifying phenomic associations of genetic variants called Pleiotropic Variability Score (PVS). While the technique could be applied to any biological entities (genes, proteins, metabolites, transcripts, microbiome, etc.), we have only tested the approach’s utility as a genomic interpretation metric. Analyses and validation of the utility beyond SNPs are out of scope for this work.

### Genotype-phenotype annotation databases

Multiple databases catalog genotype-phenotype relationships of common or rare genetic variants^56–60^. While many of these databases provide phenotype information, to the best of our knowledge only GWASdb^61^ and GWASdb v2^62^ curate phenotype codes using standard ontologies. Other databases map phenotypes defined in the original publication of genotype-phenotype discovery or replication or to generic phenotypes without an ontology to describe their relationships (e.g. OMIM phenotypes).

### Ontologies for pleiotropy assessment

An ontology is defined as a compendium of knowledge about a domain (e.g., medicine, biology) represented using a controlled, standardized vocabulary for describing both concepts and the relationships between concepts. A well-defined ontology can aid in the logical abstraction of knowledge about a field of study (e.g. gene ontology to summarize biological function of a gene or enrichment analyses of gene sets). The Open Biological and Biomedical Ontology (OBO) Foundry (See: www. http://www.obofoundry.org/) lists 136 active biological, healthcare, and clinical ontologies. Out of these, 35 ontologies incorporate phenotype concepts. Some of these ontologies are species specific (e.g., Drosophila Phenotype Ontology, Fission Yeast Phenotype Ontology, Human Phenotype Ontology), while others are organism agnostic and describe primary attributes (e.g. Measurement method ontology). We used human phenotype ontology (HPO) and human disease ontology (DO) to compute phenomic similarities as a pleiotropy measure of genetic variants. Ontological structure and various metrics of HPO and DO are provided in Table 1.

**Table 1:**
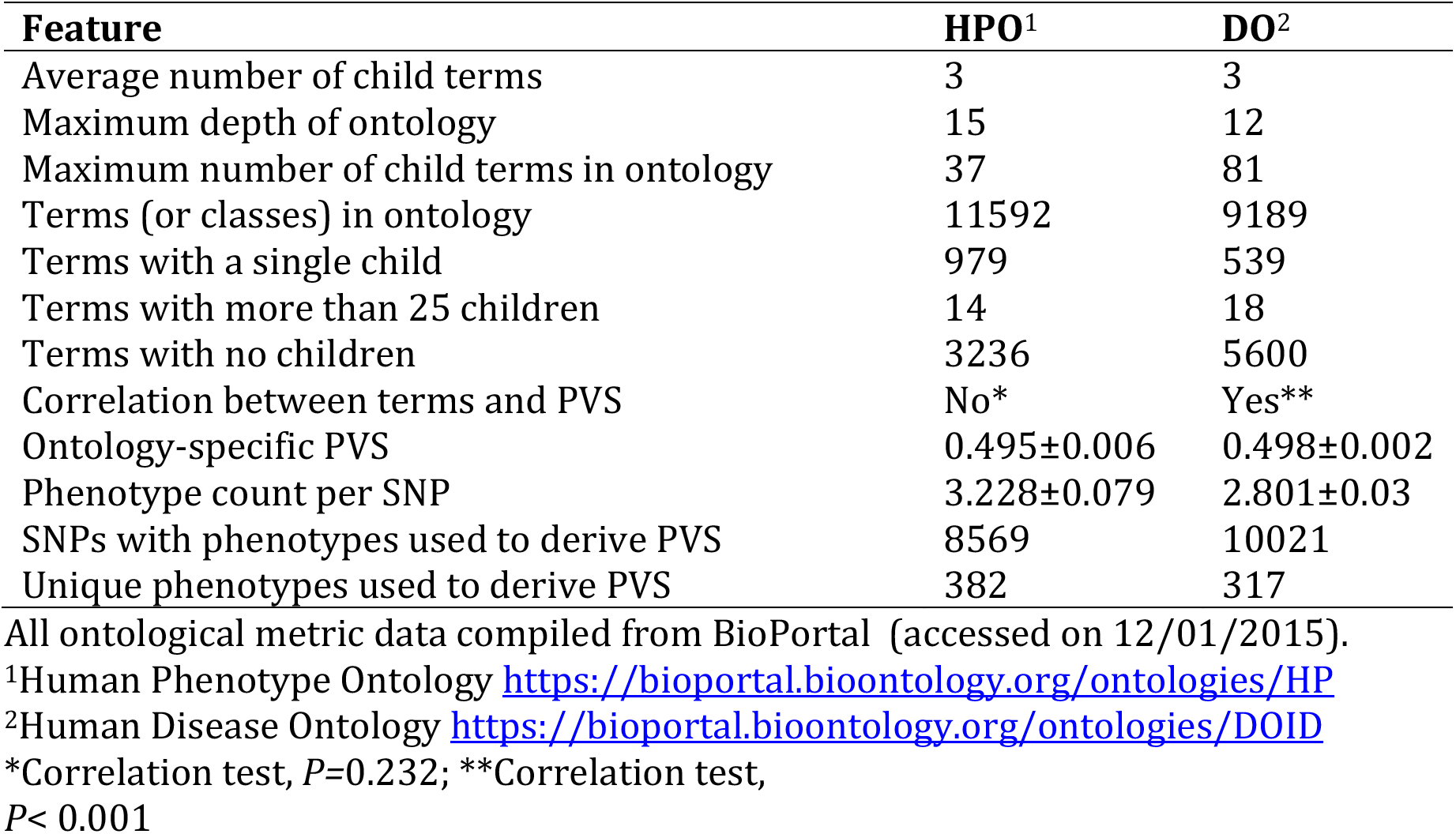
Summary of Human Phenotype Ontology and Human Disease Ontology used to derive PVS

#### Human Phenotype Ontology (HPO)

HPO is a structured and controlled vocabulary that relates the phenotypic features encountered in human hereditary and non-hereditary disease ^63, 64^. HPO has been used in comparative disease analysis tools designed for clinical phenotyping and diagnosis (e.g. Phenomizer^65^). Clinical modifier, Mode of inheritance, Mortality/Aging, and Phenotypic abnormality represent the base classes of HPO. HPO has more than 11,000 terms and over 115,000 annotations of hereditary diseases and provides annotations to approximately 4,000 common diseases. We have used the data-version/format-version: 1.2 with date stamp releases/2015-02-14 for the analyses described in this study.

#### Human Disease Ontology (DO)

DO is a hierarchical, controlled vocabulary for classifying human disease ^66, 67^. The base classes of DO are: disease by infectious agent, disease of anatomical entity, disease of cellular proliferation, disease of mental health, disease of metabolism, genetic disease, physical disorder, and syndrome. DO has been used as the knowledgebase for disease enrichment analyses^68^. We have used the data-version: releases/2015-02-12 with date stamp 11/02/2015 for the analyses described in this study.

### Computing phenomic similarity of clinical traits and human diseases

The pleiotropic score of a variant is derived using semantic similarity metrics computed using HPO and DO. Semantic similarity metrics compute an objective distance between terms in a knowledgebase or ontology, where the distance is based on the meaning or semantic content as opposed to similarity which can be extracted from their syntactical representations. Multiple methods have been implemented to derive semantic similarity scores that use different features (topological similarity, natural language processing, text mining, or statistical similarity) of knowledge corpus^69^. Semantic similarity can be derived using a diverse set of enumeration methods. Some of the commonly used metrics in biological and clinical domain include the methods proposed by Resnik^70^, Lin^71^, and Wang et al.^72^. Semantic similarity scores are typically computed on a pair-wise basis or set-wise; furthermore, these scores can be summarized as min, max or best metric average (BMA). These metrics have been part of software frameworks to calculate gene-function or gene-disease similarities and similarities of terms in ontologies^52, 65, 73, 74^. We used an ontology agnostic software framework (Semantic Measures Library and Toolkit, http://www.semantic-measures-library.org/) to compute phenomic similarities as part of PVS algorithm and derive the best-representative scores from the library of 128 methods. To avoid known methodical bias observed in phenotype similarity associated with diseases^75^ and to address the coverage of phenotype ontologies^76^, we performed a data-driven comparative analyses of various semantic similarity measurements to identify the most robust metrics to derive pleiotropic similarities across phenotype ontologies.

#### Phenomic similarity of genetic variants using Resnik method

The semantic similarity of a variant with two phenotypes (*t_1_* and *t_2_*) can be computed using Resnik method^70^ by deriving the similarity between terms in the context of an ontology (HPO or DO) using information content (IC) of their most informative common ancestor (MICA) of two terms (*t_1_* and *t_2_*) using the following equation:

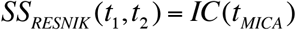

#### Phenomic similarity of genetic variants using Stojanovic method

The Stojanovic method is a graph theory based method that uses a shortest path algorithm. The similarity between two terms is computed by considering their relative location in a common hierarchy, specifically the shortest path to reach a common term. The Stojanovic method does not need to traverse through the entire ontology to derive the similarity but the computation will terminate upon finding a common parent term using shortest path.

#### Group wise integration of pairwise scores

Pairwise similarity metrics like Resnik and Stojanovic can be summarized as (*SS_Max_*), minimum (*SS_Min_*), mean (*SS_Mean_*), and best-match average (*SS_BMA_*). If a variant is associated with two phenotypes, similarity is computed in pair-wise manner; for more than two phenotypes, a group wise integration is used. Each of the group wise semantic similarity computing methods is briefly discussed below and a more detailed account can be found in the literature.

#### Maximum similarity of pairwise scores

The *SS_Max_* metric computes the maximum semantic similarity score over all pair wise similarities (*S_ij_*) using the equation:

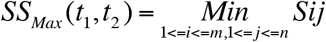

#### Minimum similarity of phenotypes

The *SS_Min_* represents the minimum semantic similarity score of all pairwise scores (*S_ij_*) using the equation:

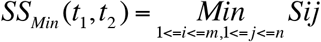

#### Average phenomic similarities

The *SS_MEAN_* computes the average semantic similarity of all pairs of term sets in respective ontologies using the equation:

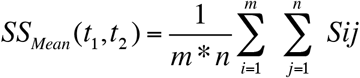

#### Best-match average of phenomic similarity

The *SS_BMA_* method derives a best-match average from the scores using the following equation. BMA is obtained from the similarity matrix S with by computing the average of all maximum similarities on each row and column.

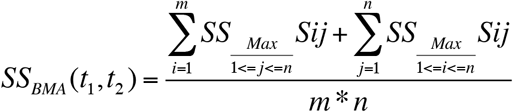

We generated 172 combinatorial group similarity metrics by implementing each possible dependency combination as in Fig S1. We tested several metrics from each fundamentally different basis of evaluating the ontology in combinatory fashion, including 1) single node evaluation of intrinsic information content measures (3 methods:), 2) similarity of pairs of 2 nodes (3 methods based on node set:, 3 based on node information content:, and 2 based on edges:), 3) group similarity of measures of pairs (5 methods:), and 4) group similarity of measures of many individual nodes (2 methods based on connectivity:, 2 methods based on information content:). Semantic similarity of pair of diseases were computed using the Java-based Semantic Measures Library and Toolkit^77^.

### Pleiotropic Variability Score

Multiple combinations of semantic similarity metrics and approximation yields an extensive library of scores. Integrating these scores into a single parameter is ideal to incorporate it in genome analyses and interpretation workflow. To understand how each of these metric influence scoring, we compiled data on various metrics across both ontologies (See Figure S1-S5). After assessing 128 semantic similarity method combinations, we selected the top six methods based on consensus correlation and scale of assessment (i.e. local similarity vs. global similarity of two terms or concepts) to define pleiotropy. After computing the semantic similarity using these methods, we defined PVS as the inverse of the semantic similarity.

Pleiotropy Variability Score using HPO was derived using the following equation:

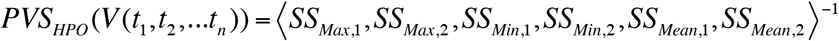

Pleiotropy using DO was derived using the following equation

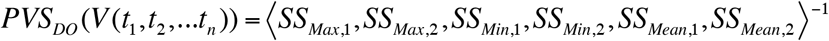

Where top2MinMetrics were MIN_Stojanovic and MIN_Sim, top2MaxMetrics were MAX_Resnik and MAX_Sim, and top2MeanMetrics were AVERAGE and BMA. Finally, we average the score across both ontologies for variants with both phenotypes to offer a complete spectrum of disease and phenotypes represented in the equation using the equations:

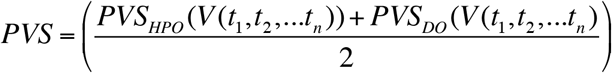

Different semantic similarity metrics are characterizing set similarity of terms with different scopes of consideration. To obtain a better coverage across all similarity scores that reflect the full range of similarity scopes we use an average of the top pair of metrics (Max, Min and Mean) from each of the fundamentally different scoring approaches. By incorporating multiple methods, PVS incorporates multiple layers of information in a single score. For example: *Min* metrics provides the intrinsic and extrinsic similarity of terms using global similarity trends in ontology. *Max* based metric provides information content-based similarity using local similarity. *Mean* or *BMA* based method capture shared all the best matching semantic similarity values for each term. To conclude PVS provides a single metric by wrapping the entire compendium of scoring methods to capture phenomic similarity to quantify pleiotropy.

### Biological implications Comparison with other variant interpretation scores and methods

Variants were mapped to genes and stratified by PVS score levels (L1, L2, L3, L4) and tested for biological and functional enrichment with Reactome pathways and Gene Ontology (molecular function) annotations using Enrichr (https://maayanlab.cloud/Enrichr/). We compared PVS with various genome interpretation metrics including FATHMM^78^, SIFT^79^, PolyPhen-2^80^, CADD^81^, GERP++^82^, LRT^82^, MetaLR, MetaSVM^83^, MutationAssessor^84, 85^, MutationTaster^86^, PROVEAN^87^, PhastCons^88^, PhyloP^82^, SiPhy^89^, CONDEL^90^, and VEST3^91^. Variant annotations were obtained from dbNSFP^92^ and Ensemble Variant Effect Predictor (VEP)^93^ and all analyses were based on the impact on the canonical transcript. We have also compared the PVS of variants by SNP location, CNV annotation to understand the biological implications of PVS. For both analyses SNP position was mapped to SNP type and CNVs using SNP-Nexus pipeline.

### Clinical validation

Canonical annotations from biomedical genomics database can be used for developing theoretical models and prediction methods. To test the validity of PVS in genome interpretation, we tested the incidence of pleiotropic variants using an EMR-linked genomic biorepository. Our goal is to determine the disease comorbidity profile within pleiotropic allele carriers in a patient population. We estimated the prevalence of pleiotropic hub carriers using our validation cohort using disease phenotype mapping and SNP based overlap of PVS scores.

### PVSAnnotate

We developed a software pipeline called PVSAnnotate (Figure 3) to incorporate the computation of PVS as part of genome interpretation. VCF data handling routines in PVSAnnotate is developed using R, and the software workflow is a shell script-based implementation amenable to integration with existing genome analyzers and interpretation packages. The pipeline uses SMLTK to compute semantic similarity scores for variants in VCF to derive the allele-specific pleiotropic score and will output a formatted VCF incorporating PVS.

## RESULTS

We designed, developed and validated a new objective scoring method for genome interpretation that will rank SNPs based on known phenotype associations. We used PVS algorithm and annotated ~12K variants in human genome. A new schema for quantifying pleiotropy is introduced and examined the unique biological functions across pleiotropic levels (L1 to L4). Our results suggests that pleiotropic levels have implicit biological meaning and comparable to scores like SIFT. PVSAnnotate, a software framework based on PVS is used to annotate personalized pleiotropy of a reference genome and population scale, genomic data from EHR-linked biobank We envisage that PVS will be part of genome interpretation pipelines and contribute to the understanding of molecular basis of disease comorbidities and network architecture of pleiotropy in human disease physiology. We demonstrate possible applications of this method by annotating a sample genome and estimating pleiotropy from an EMR-linked genomic bank using the PVSAnnotate software framework as well.

### Summary of genotype-phenotype dataset

We compiled and analyzed a total of 12, 541 SNPs with rs* identifiers and 699 phenotypes to derive canonical PVS. 72.9% of the SNPs were Transitions (changes from A <-> G and C <-> T) and 27.1% were Transversions (changes from A <-> C, A <-> T, G <-> C or G <-> T). Transitions are expected to occur twice as frequently as transitions; our dataset is indicative and thus present a good representative sample. Scores were generated with all genetic variants reported in the reference database (union of GWAS Catalog + GWASdb v2 (version gwasdb_20140908_snp_trait) + GRASP) with more than one DO (*n*=317 unique phenotypes; 88.536±22.277) or HPO (*n*=382 individual phenotypes; 72.48±13.628) associated phenotypes. Within the DO database, type 2 diabetes mellitus, coronary artery disease, myocardial infarction, cardiovascular system disease, and bipolar disorder were the most represented diseases, comprising 3.5% of all terms annotations (*n*=8,939 terms in the background). For HPO, phenotypic abnormality, Type 2 diabetes mellitus, bipolar affective disorder, myocardial infarction, and schizophrenia were most represented phenotypes, comprising 3.45% of all HPO terms (*n*=11,065 terms in the background).

### Deriving the PVS metric to quantify pleiotropy

We tested several metrics (Figure S1) from each categories in combinatorial fashion, including 1) single node evaluation of intrinsic information content measures (3 methods:), 2) similarity of pairs of 2 nodes (3 methods based on node set: 3 based on node information content: and 2 based on edges:), 3) group similarity of measures of pairs (5 methods:), and 4) group similarity of measures of many individual nodes (2 methods based on connectivity: 2 methods based on information content). We list brief descriptions of individual methods in Table 2 and outline each combination of methods assessed to find best-representative metrics to derive PVS in Supplementary Data (PVS_SS_Metrics_Tested.xlsx). We tested a total of 172 semantic similarity measurements and compiled the pleiotropy score using 6 metrics. Finally, we detail correlation consensus of 172 metrics assessed used to derive final set of semantic measures from different phenotype lengths (See Supplementary Data PVS-Phenotype_Len_Correlations.xlsx). Correlation plots of methods were examined, and we picked methods that agree across both ontologies using R^2^ as a parameter. Finally, we picked a total of 6 methods that provides depth, breadth and reduce the bias to top terms. It should be noted that semantic similarity algorithms are sensitive to specificity of information and severity of impact increases with more general phenotypes (Figure S2-S5).

**Table 2:**
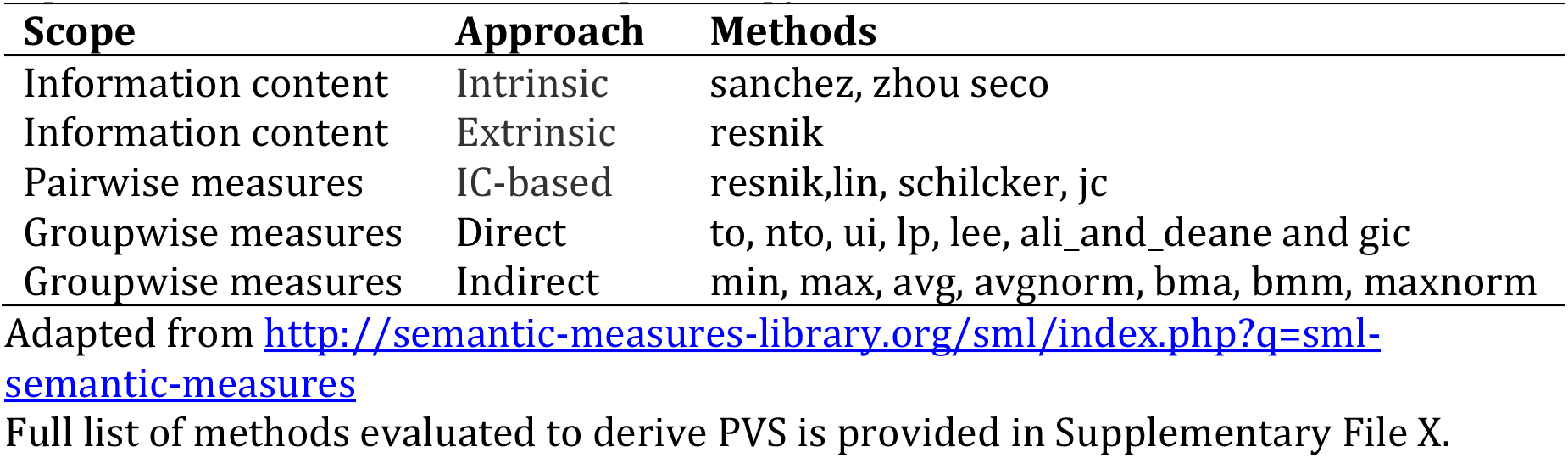
Different semantic similarity measures assessed to identify best-representative methods to derive pleiotropy

### Quantifying pleiotropy using PVS

We provide a flowchart summarizing the PVS algorithm in Figure 1 along with the semantic similarity scores computed using phenotypes associated with 12,646 variants. Additionally, we detail the PVS derived from the semantic scores of the metrics computed using HPO, DO, and the final integrated score within the same figure. We calculated quartiles of the PVS (Kolmogorov-Smirnov, *P*<0.01) and organized the scores into four different levels of pleiotropy as follows. We hope the levels of pleiotropy using a simple L1-to-L4 system would further help to establish the use of PVS as an easy to interpret metric as part of genome interpretation workflows.

- L1 (no or low pleiotropy; 0.35±0.09), a variant with a PVS score *x* for a variant *V* is considered L1 when *Vx*{∈ *L*1|0.264 ≤ *PVS* < 0.436}; An example for an L1 PVS SNP would be rs1002552 with a PVS=0.308; PVS-Class=L1; associated phenotypes used for estimating pleiotropic variability are acute lymphocytic leukemia and acute leukemia.
- L2 (low or medium pleiotropy; 0.48±0.04) when *Vx*{∈ *L*2|0.436 ≤ *PVS* < 0.52}; An example for an L2 PVS SNP would be rs1004446 with a PVS=0.51; PVS-Class=L2; associated phenotypes used for estimating pleiotropic variability are type 1 diabetes mellitus and multiple sclerosis.
- L3 (medium or high pleiotropy; 0.54±.02) when *Vx*{∈ *L*3|0.52 ≤ *PVS* < 0.56}; An example for an L4 PVS SNP would be rs1046747 with a PVS=0.543; PVS-Class=L3; associated phenotypes used for estimating pleiotropic variability are asthma, leukemia, lymphoma, lymphopenia, cardiovascular system disease
- L4 (high or super pleiotropy; 0.565) when *Vx*{∈ *L*4|0.56 ≤ *PVS* < 0.673}; An example for an L4 PVS SNP would be rs964184 with a PVS=0.565; PVS-Class=L4; with more than 45 disease, phenotypes, or traits associations.

**Figure 1:**
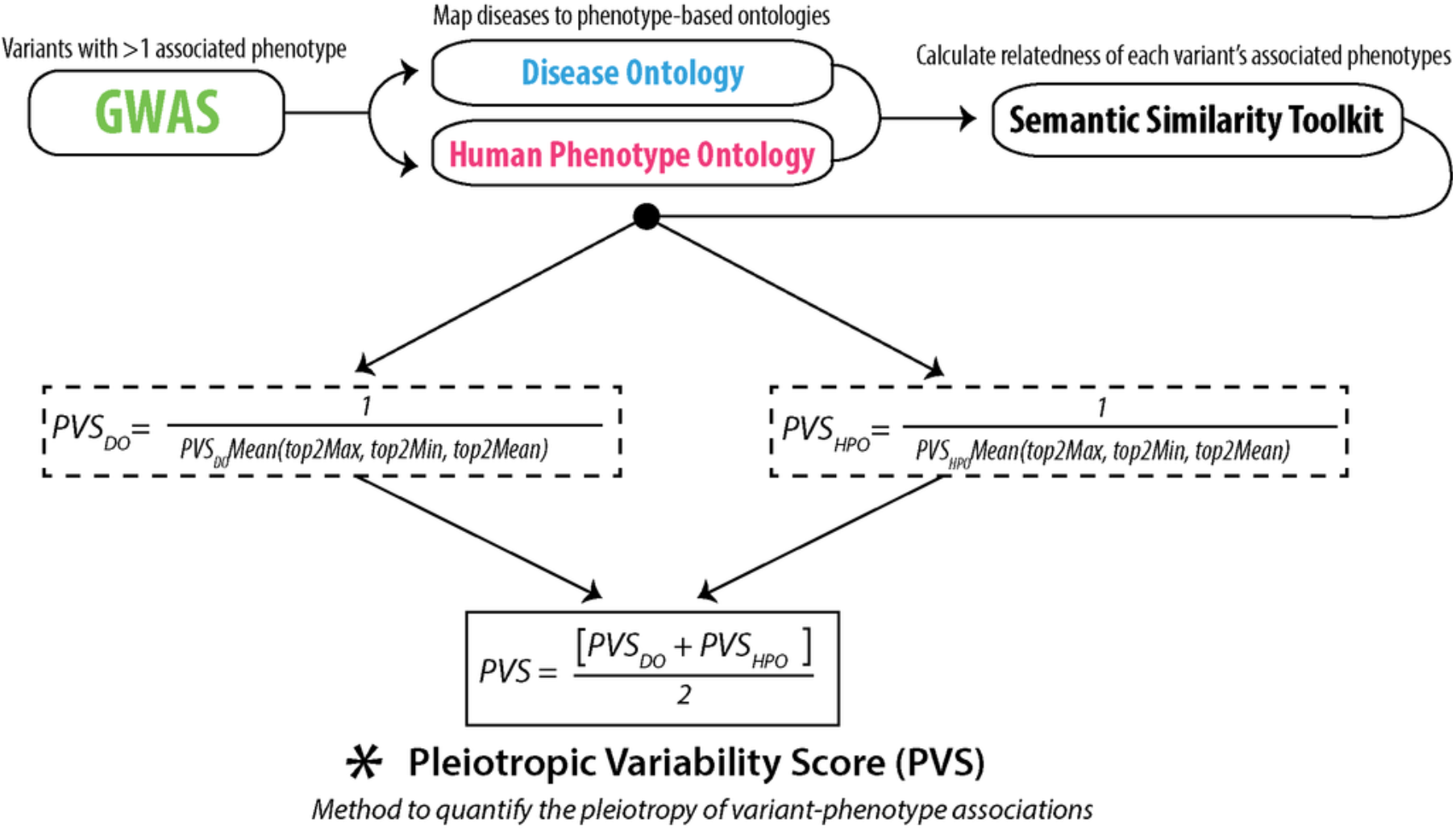
Graphical outline of PVS methodology

#### Determining disease prevalence for patients with highly pleiotropic variants

We sought to validate the PVS metric in a clinical cohort by first identifying any pleiotropic variant carriers and then assessing prevalence of associated disease phenotypes within these carriers. Specifically, we determined for each hub, the amount of individuals that had any combination of diseases associated with the hub. For each rsID-allele combination we identified total number of patients with the risk allele (numwithallele), number of patients heterozygous for the risk allele (numheterozygous), number of patients with a homozygous alternative genotype (numhomoalt). We then compared the disease frequency data including total number of diseases associated with variant (diseases) followed by number of patients with all but one disease in decrement order (d, d1, d2…d10) between groups (See Figures 2, 3 and Supplementary Data File: PVS-BioME-Validation_Results.xlsx).

**Figure 2:**
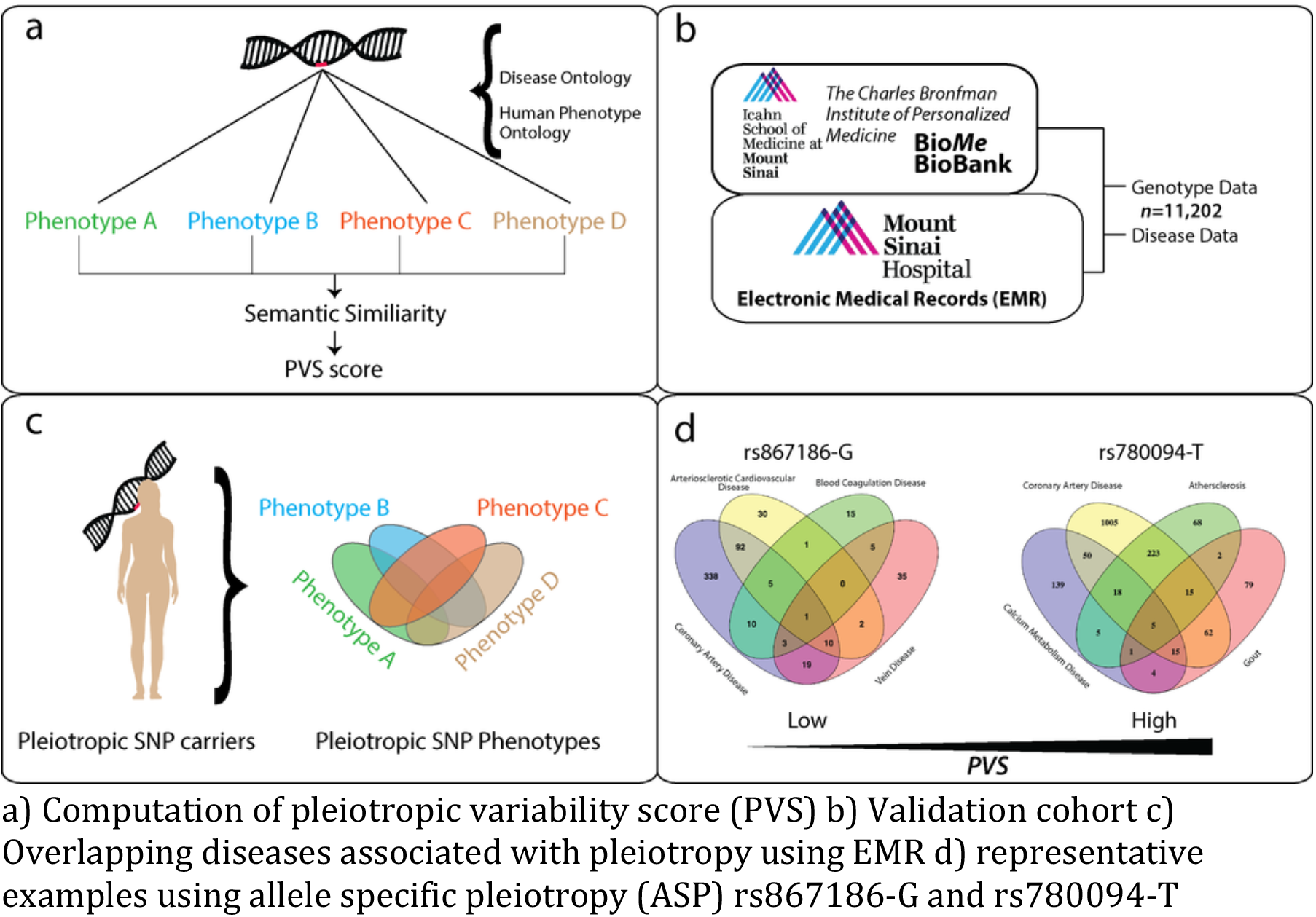
Visual graphic of validation approach used for Pleiotropic Variability Score using an electronic medical record (EMR) linked biobank (BioME)

**Figure 3:**
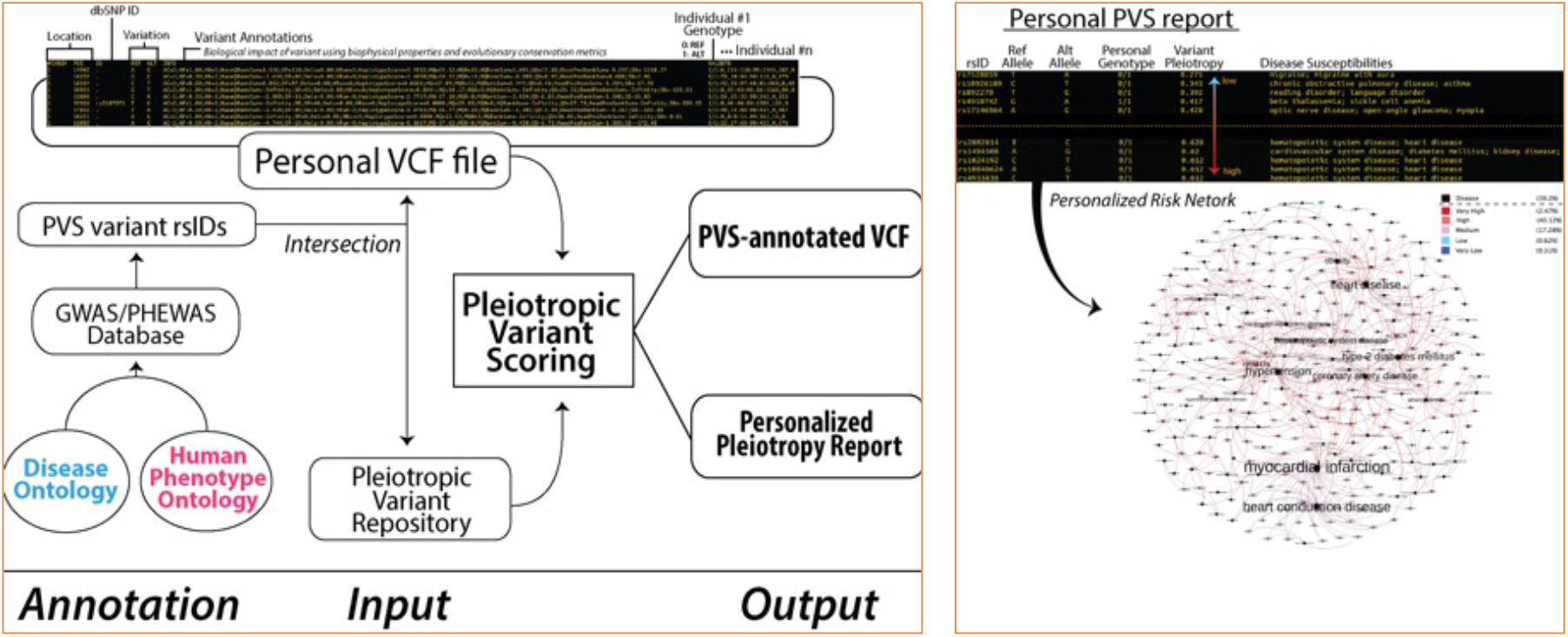
Flow-chart of software pipeline (PVSAnnotate) to incorporate pleiotropic variability as part of genome annotation and interpretation and visual graphic of a personal PVS report

### Comparing PVS with canonical annotations of genetic variants

We mapped 12,541 variants to 4,053 genes, 15,383 transcripts, and 1,874 regulatory features. These SNPs affect 33 different biotypes including regions (e.g. enhancers), transcription factor binding sites, and non-sense mediated decay. We outline a summary of variation consequences for the subset of coding variants along with the variant annotations compiled using Ensembl VEP and dbNSFP databases (See Figures 4, 5, 6 Supplementary Data: PVS-SNPs-Annotation.xlsx).

**Figure 4:**
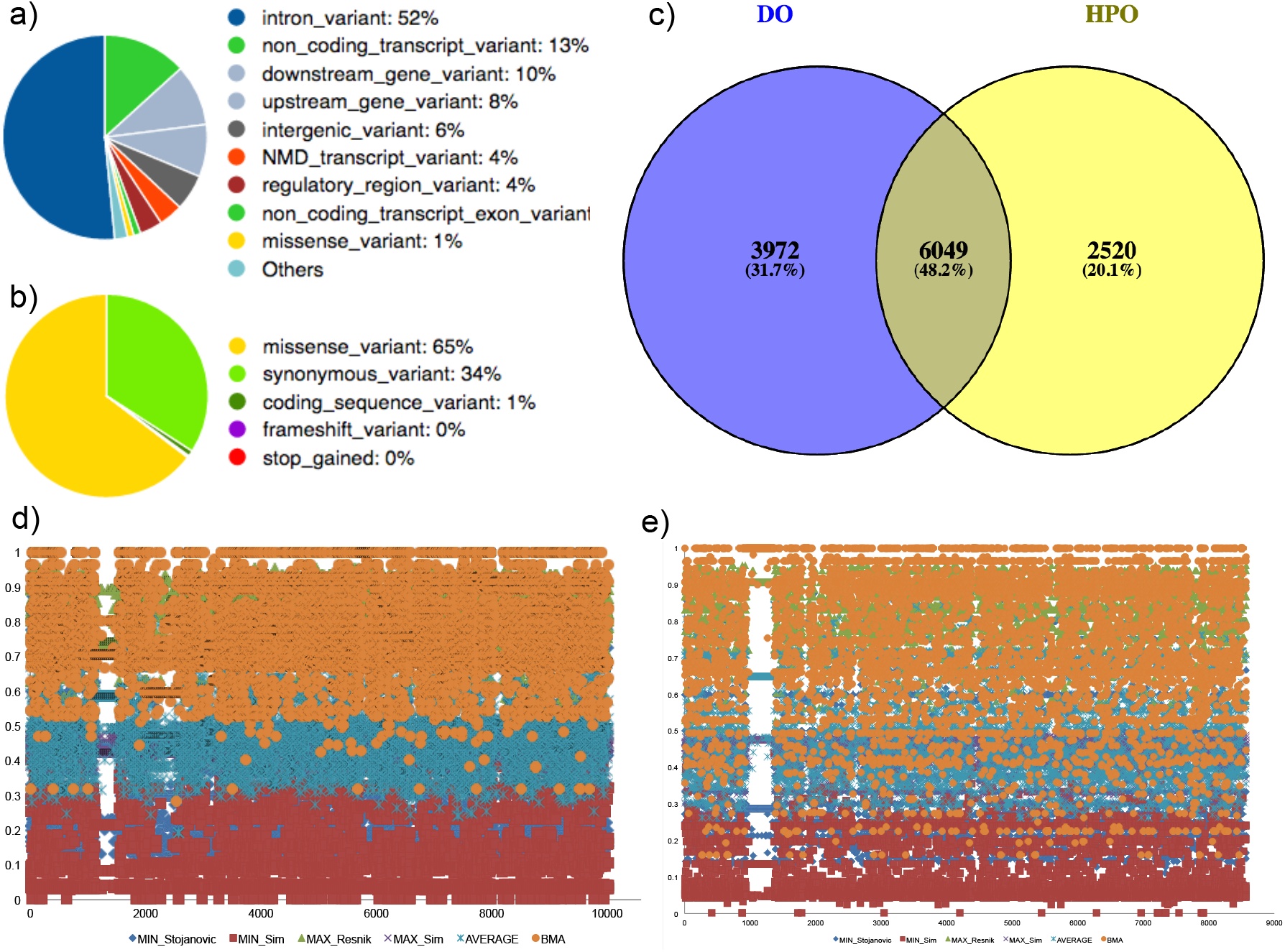
Summary of variants and individual scores used in PVS. Variant consequences a) all and b) coding sequence), c) SNP-phenotype coverage and semantic similarity scores using six metrics using e) DO and f) HPO ontologies

**Figure 5:**
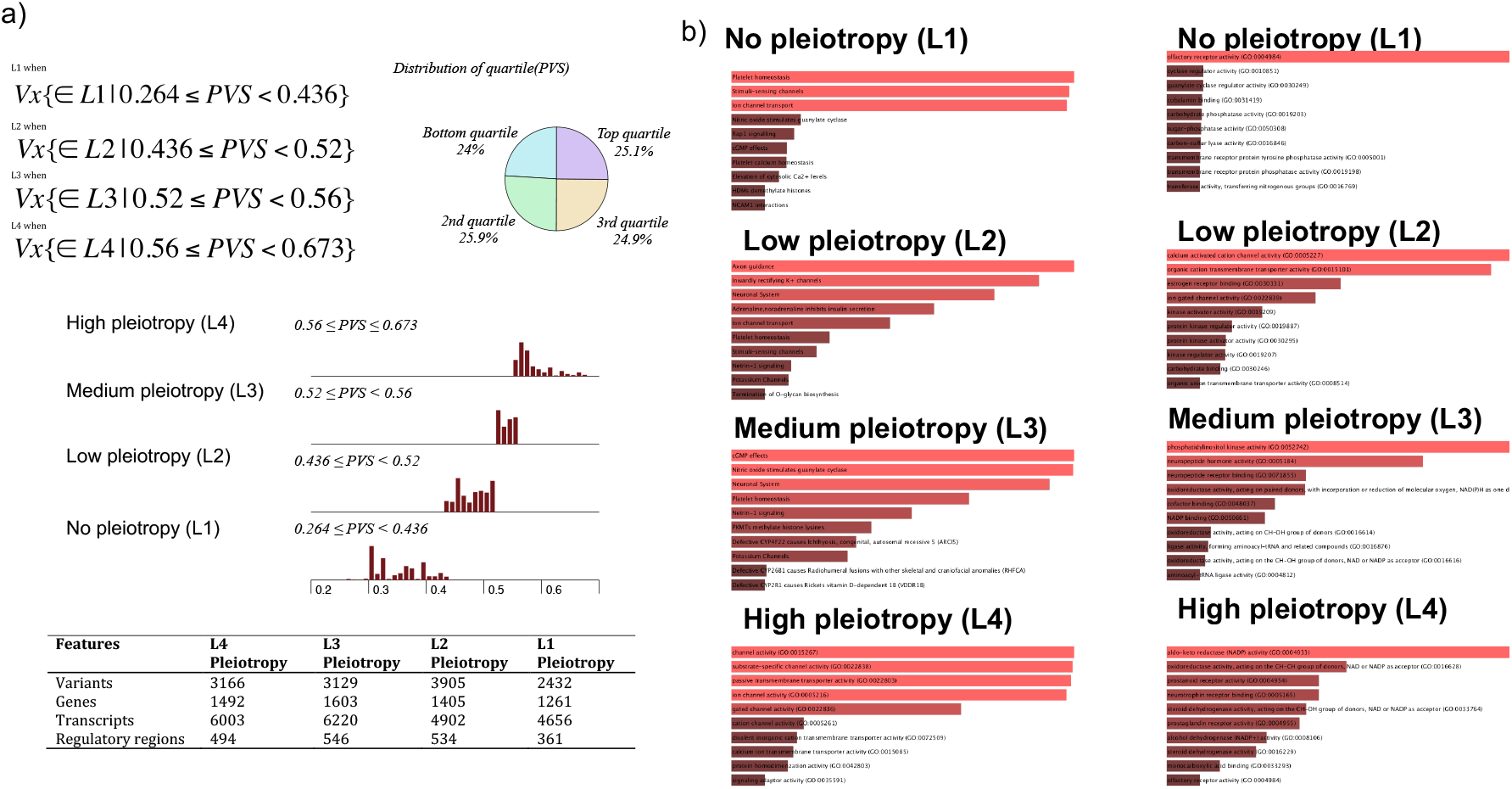
Pleiotropic variability scores of genetic variants a) distribution of scores estimated b) Functional enrichment of genes mapped from the variants (Reactome pathways and Ontology: Molecular Functions) across different levels of pleiotropy

**Figure 6:**
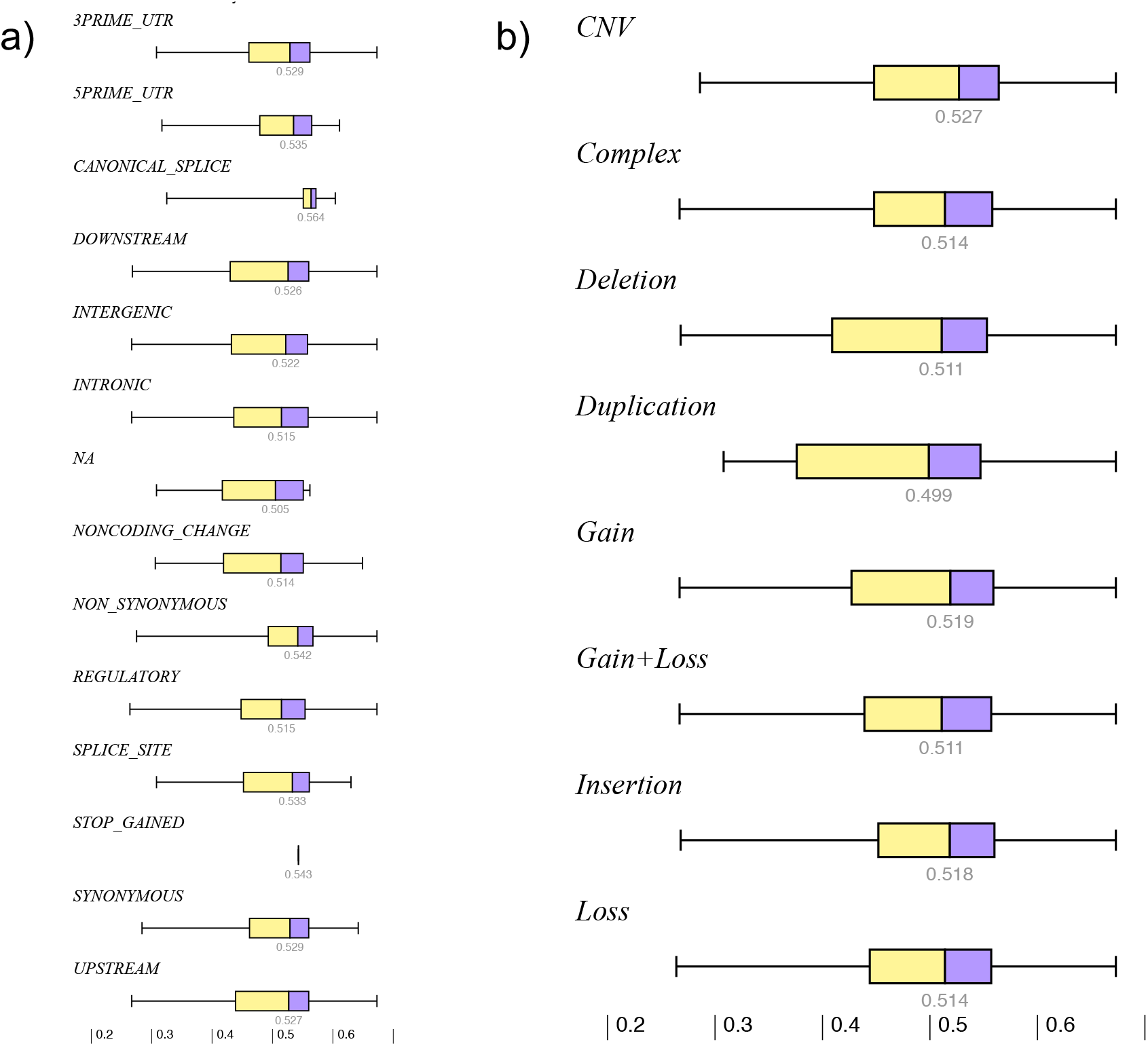
Distribution of PVS across SNP types and copy number variations

#### Allele specificpleiotropy of rs867186-G

The variant rs867186 (1KG-MAF: 0.0001) is a coding variant encoded in protein c receptor (*PROCR*) gene. *PROCR* (synonyms *CCCA, EPCR*, or *CCD41*) is a serine protease and involved in blood coagulation pathways, malaria, and cancer^96^. Variants in this gene have been associated with myocardial infarction, venous thromboembolism, late fetal loss during pregnancy and play a key role in inflammatory pathways ^97, 98^. The variant is associated with seven phenotypes that include traits (protein C levels, coagulation factor levels, anticoagulant levels, hemostatic factors and hematological phenotypes, and D-dimer levels) and diseases (coronary heart diseases and coronary artery disease) across three populations (EUR, AFR, and SAN). These phenotypes were mapped to four terms in ontologies (coronary artery disease, arteriosclerotic cardiovascular disease, blood coagulation disease, and vein disease). Out of 1985 patients with the rs867186-G allele (5.7% homozygous patients) only 115 patients had homozygous alleles and 478 patients had one or more of the associated diseases (24.0%).

#### Allele specific pleiotropy of rs780094-T

The rs780094-T variant is a coding SNP on glucokinase (hexokinase 4) regulator (*GCKR*) gene and inhibits glucokinase in the liver and pancreas ^99, 100^. This variant is associated with multiple clinical traits including LDL cholesterol, triglycerides, and phospholipid levels, as well as and disease phenotypes such as calcium metabolism disease, coronary artery disease, atherosclerosis, and gout across five different populations (EUR, AFR, SAN, ASN, and AMR). Within our population, out of the 5,442 patients with the risk allele (20% homozygous patients), 25.6% individuals had one of the four diseases. Collectively these examples show the prevalence of genotypes with multiple phenotypes is lower than expected. This trend could be driven by the age of the individuals, ethnic origin, individualized susceptibility profiles, environment, and the influence of modifiable risk attributes. Replicating our initial findings using a large genotype-phenotype database will provides an estimate of pleiotropy on a population-scale and could help in identifying trends in adaptive evolution and antagonistic pleiotropic trends.

### Biological Implications and Comparison with variant interpretations metrics

To understand the biological role of the genes harboring SNP, we have conducted enrichment analyses using genes mapped to the different levels. Some of the key Reactome pathways enriched across PVS levels includes platelet homeostasis (L1); axon guidance (L2); cGMP effects (L3) and channel activity (L4). Similarly, molecular function terms enriched across PVS levels includes olfactory receptor activity (L1); calcium activated cation channel activity (L2); phosphatidylinositol kinase activity (L3); and aldo-keto reductase (NADP) activity (L4).

By mapping the distribution of PVS by SNP types and CNV types (Figure 6), interesting biological insights can be gained (Kruskal-Wallis, p < 0.001; unequal medians; Figure 6). For example, SNP types mapped “canonical splice site” have a mean PVS of 0.564. This is an interesting finding as splicing events are considered as an important mechanism by which genes like G-protein coupled follitropin receptor, SR proteins and TMPRSS2/ERG fusion transcripts have demonstrated pleiotropic properties. Further analyses and function test experiments may yield novel biological insights on how low or higher order pleiotropy computed by PVS may influence biological mechanisms.

We have compared PVS with several genetic variant interpretation metrics including SIFT, PolyPhen2, etc. to illustrate its utility as a genomic interpretation tool. PVS and SIFT have a negative correlation (Correlation test, p < 0.001, *n*=481 SNPs; Figure S6). From a biological perspective, this correlation is potentially indicative of higher the pleiotropy; lower the damage estimated by SIFT approach. We did not observe a correlation for PolyPhen2 scores and PVS (Correlation test, p = 0.669, *n*=557). Further analyses using expanded set of SNP data and stratified analyses by levels may yield correlations.

### PVSAnnotate - A software framework for quantifying pleiotropy of genomes

The PVSAnnotate software pipeline and all related source code and data are available in the public domain. Source code and data to implement software pipeline (PVSAnnotate; See Figure 3) to annotate genomic data using pleiotropic variability score is available from the URL: https://bitbucket.org/dudleylab/pvs. The potential users should note that our knowledge of genotype-phenotype associations is rapidly expanding, hence confirming data provenance is critical to ensure the reproductivity of these scores.

### Annotating a genome using PVSAnnotate

The PVSAnnotate software pipeline can import a standard Variant Calling Format (VCF file) and will annotate all variants with PVS scores. The variants were segregated into individuals and mapped to the genotype-phenotype table and associations will be gathered. The pipeline can also produce personal PVS report and personalized risk network as by integrating with various popular bioinformatics tools including Cytoscape (See Figures 2 and 3). To illustrate the application of PVSAnnotate, we have annotated variants from whole genome profiling of a patient (NA12878, genome in a bottle sample) and provide the output (See Supplementary Data File: PVSAnnotate-VCF-Output.xlsx). For this example, we obtained the individual genome in VCF format with SNPs and genotype from ftp://ussd-ftp.illumina.com/hg19/8.0.1/NA12878. We mapped the SNPs to PVS catalog to EBI-GWAS catalog to check the risk alleles (https://www.genome.gov/admin/gwascatalog.txt) by assuming an EA ancestry. Risk alleles were then mapped to phenotypes using canonical annotations in PVSAnnotate with GWASdb v2(http://jjwanglab.org/gwasdb) as a bridge. There is no phenotype data was available for this individual at the time of our analyses in the public domain other than a pharmacogenomic variant information (http://www.nist.gov/mml/bbd/ppgenomeinabottle2.cfm). Our annotations provide a pleiotropic assessment of various variants.

## Discussion

Pleiotropy - a fundamental biological phenomenon discovered in the ‘50s where a single genomic feature affects multiple phenotypes has growing relevance in the current era of precision medicine and high-throughput biology. With the availability of cost-effective phenotyping algorithms, affordable genome profiling, and digital health tools, such as EMR-linked biobanks, more genotype-phenotype associations have been discovered in the last decade than any time in the history of biology and medicine. We can now leverage genome-scale pleiotropy for various applications, including genome interpretations and complex disease risk estimations. We designed the PVS system to make use of genotype-phenotype annotations and ontologies for systematic and reproducible computation of phenotype pleiotropy of genetic variants.

PVS encompasses the pleiotropic variability of a variant by computing the similarity or diversity of all phenotypes associated with a genetic variant. Using a multitude of semantic similarity metrics, PVS can be utilized in multiple ways, such as finding novel variants from a WGS or WES study, or to assess if a gene or variant is part of a known phenomic hub. A genetic variant with high PVS can help to reveal the biological mechanisms underlying all associated phenotypes. PVS is a metric that illustrates the integration of massive amount of heterogeneous genomic and phenomic data could aid in developing better metrics for genome interpretations. Pleiotropy, along with variations such as antagonistic pleiotropy, offer options to understand the pervasiveness of certain disease pairs and absence of others driven by genetic variants. PVS could also be used as a metric to find potential antagonistic pleiotropic trends, where the associated phenotypes do not co-manifest, on a population scale. Additionally, using PVS to systematically analyze antagonistic pleiotropy by identifying inverse comorbidity relationships in the patient population could help to identify new drug targets (pleiotropic hubs in human phenome) that drive disease pathogenesis mechanisms and could aid in developing precision therapies. Phenomic pleiotropy could also help and guide to understand molecular underpinnings of drug development strategies including drug repositioning. For example, an L1 PVS SNP rs11360 is associated with stroke; where as an L4 PVS rs877098 is associated with alcohol dependence^94^ and multiple complex diseases (bipolar disorder, coronary artery disease, Crohn’s disease, rheumatoid arthritis, type 1 diabetes and type 2 diabetes)^95^. Exploring these genomic targets may yield potential mechanistic insights and putative drug targets but the first could be target for a specific disease; whereas the second could be a pleiotropic target that could modulate multiple diseases.

In our dataset 10,021 SNPs have >= 2 DO associations and 8,569 SNPs have >=2 HP associations. Top 3 phenotypes with SNP associations from DO were type 2 diabetes mellitus (*n*=1471), coronary artery disease (*n*=1288) and myocardial infarction (*n*=1231). Top 3 phenotypes with SNP associations from HPO were phenotypic abnormality (*n*=1144), type 2 diabetes mellitus (*n*=958), and bipolar affective disorder (*n*=705). These examples indicates that phenotype-based data also have inherent differences in genome-phenome association and hence there is a need to bring evidence from different annotations and ontologies. In this work, we decided on a combination of semantic similarity metrics as the most appropriate method for estimating pleiotropy to overcome score inflation biases due to overrepresentation of phenotypes within a single class. Specifically, we used products of two metrics for each approximation (*Max, Min and Average*), deriving PVS from Max similarity would compute the similarity represented by the absolute terms in a given ontology. Incorporating all three approximations into PVS creates a more robust metric, which overcomes limitations of each approximation individually. For example, if we consider two genetic variants (VarA and VarB): VarA is associated with two different phenotypes (infection and cancer) and VarB is associated with the pair of clinically different but biologically related conditions (Parkinson’s disease and Alzheimer’s disease). During similarity computation *Min* based approximation rank both of these variants with high pleiotropy and lead to scoring of any two terms from same or different part of ontology as pleiotropy. *Max* would rank only the extreme example as pleiotropic, but do not account for clinically interesting differential diagnoses (Alzheimer’s disease and Parkinson’s Disease). The *Average* method assigns the variant associated with infection and cancer as highly pleiotropic (infection and cancer) but the other variant will get an estimate on the medium scale of pleiotropy.

Previous attempts to classify or score pleiotropic effects^101–104^ including seven types of pleiotropy, Fisher’s geometric model, and degree of pleiotropy rely on manual assignment of phenotypes or traits to a gene, which is difficult to scale and keep up-to-date. PVS applies a more pragmatic approach by using algorithms to consider the entire, current phenotype knowledgebase. Such approaches are becoming increasingly complex with the rapid growth of low-cost, high-depth genomics projects^9, 27, 28, 105, 106^. PVS offers unbiased estimation of pleiotropy using semantic similarity as a surrogate measure for deriving pleiotropy using multiple ontologies. Compared to a previous method that relies on counts of phenotypes and a degree of surprise scoring^107^, our method utilizes a systematic approach to quantify pleiotropy by considering the inherent relationships between clinical traits and disease phenotypes in a cohesive and automated fashion.

The automated clinical-grade genome interpretation offers many challenges to clinicians and translational bioinformatics communities. Efficient and precise analysis and interpretation of genomic data are critical for its use and for application in risk estimation of complex diseases, diagnoses of rare disease, pharmacogenomics, and tumor profiling for personalized, cancer-specific drug recommendation. As pleiotropy is a crucial mechanism that drives multiple diseases and phenotypes pathogeneses, direct quantification of pleiotropy is a critical annotation layer that should be incorporated as part of genome interpretation workflows. PVS and PVSAnnotate collectively fill this gap by providing a method and software for incorporating pleiotropic variability as part of genome analyses.

### Limitations and future work

In this work, we designed, developed, and implemented a new computational approach to quantify pleiotropy of phenotypes associated with genetic variants. Our work has several limitations. In the current implementation of PVS, we calculate the semantic similarity-based estimates on two ontologies. Using ontology merging methods and incorporating additional ontologies (e.g., Experimental Factor Ontology) could improve the scoring system and will be incorporated into future versions of the PVS. Additionally, some of the genotype-phenotype databases (GWAS Central, dbGAP) provide mapping to Medical Subject Headings (MeSH). Computing semantic similarity score on the entire MeSH tree will be computationally expensive but could be useful. The broad nature of the MeSH terms could squeeze the phenotype similarity scoring to extremes when deriving *Min, Max*,or *Average*, but the addition and merging of additional ontologies would help in keeping the PVSAnnotate pipeline up to date, as well as making it more robust.

Incorporating other biological data types, like expressed quantitative trait loci (eQTL) data^108^, gene-tissue annotations^109^, and gene-gene interactions^109^ for refining genotype-phenotype annotations could further improve the scoring and quantification of pleiotropy. PVSAnnotate only performs the computation of the terms that are compiled in the database and this mapping is performed using *rs* identifiers, which are not stable. As such, future version of PVSAnnotate will use one or more identification system or mapping strategy to connect variants to phenotypes. Ontologies are often merged, revised, or deprecated, which could affect stability of ontological terms and could influence the scoring system. It should also be noted that different classification schemas have been proposed to describe pleiotropy in biology (for example, Hodgkin proposed a seven-type classification schema that includes artefactual, secondary, adoptive, parsimonious, opportunistic, combinatorial, and unifying. In the current implementation of PVS do not account these differences.

## Conclusion

Current generation of genomic variant scoring algorithms and interpretation tools often use genomic conservation, evolutionary substitution and other cross-species alignment driven approaches to rank and prioritize genomic variants. These scoring systems have been used as utility to rank variants for clinical use of genomic variants. Such scores do not account the growing evidence from genome-wide association studies and phenome-wide association studies. In this work, we are introducing a new genomic variant interpretation metric PVS, developed using a rigorous quantitative assessment framework that provides a score to depict pleiotropy by accounting all phenotypes associated with a genomic variant.

We designed, developed, and validated an objective scoring method that will rank and classify variants based on their associated phenotypes. This metric, PVS, can quantify the SNP-phenotype associations through calculating semantic similarity from HPO and DO ontologies. PVS describes the pleiotropic nature of the variant of interest complementing existing genome-interpretation scores. Standard genetic variant filtering applications can use PVSAnnotate to filter variants according to pleiotropy. For example, SNPs with higher PVS (highly influential across the phenome) study phenotypic correlations or SNPs with lower PVS (low influence; novel) for explorative research. We have extrapolated the concept of PVS from SNPs to genes and defined pleiotropic hubs. From a clinical perspective, PVS helps interpret allelic heterogeneities and helps assess genetic variant associations with the same disease class (e.g., prostate cancer and neuroendocrine prostate cancer) or several distinct diseases (e.g., myocardial infarction and cancer). We envisage that PVS will further contribute to understanding the molecular basis of disease comorbidities and network architecture of pleiotropy in human physiology.

## Availability

Software and data discussed in this manuscript to compute pleiotropic variability score is available from the URL: https://bitbucket.org/dudleylab/pvs

## Supplementary Data

Supplementary Data include six figures and 6 Excel files and can be found with this article online.

## ACKNOWLEDGEMENTS

Authors would like to thank Harris Center For Precision Wellness and Icahn Institute for Genomics and Multiscale Biology (http://icahn.mssm.edu/departments-and-institutes/genomics), of Mount Sinai Health System for infrastructural support. Authors would like to acknowledge Corey Xu, Ben Readhead, Brian A. Kidd, Douglas Ruderfer, Pei Wang and Rong Chen for discussion.

## FUNDING

This work was funded by the following grants from National Institutes of Health: National Institute of Diabetes and Digestive and Kidney Diseases (NIDDK, R01-DK098242-03); National Cancer Institute (NCI, U54-CA189201-02) to JTD and National Center for Advancing Translational Sciences (NCATS, UL1TR000067) Clinical and Translational Science Award (CTSA) grant to JTD and KS. The funders had no role in study design, data collection and analysis, decision to publish, or preparation of the manuscript.

## AUTHOR CONTRIBUTIONS

KS, MAB and BSG wrote the manuscript, KS, MAB and BSG curated the data and performed the analyses, validation and developed the software pipelines. KS, MAB, BR, BAK, PW, RC contributed ideas and participated in derivation of the scoring method. JTD contributed to the overall planning of the project, scoring method and the manuscript. All authors read and approved the final manuscript.

## COMPETING AND FINANCIAL INTERESTS

KS: Received salary, consulting fees or honoraria from Philips Healthcare, Kencor Health, OccamzRazor, Alphabet, McKinsey & Company, BCG, LEK. Employee of AstraZeneca at the time of publication; MAB: none; BSG: None declared; KWJ: None declared. JTD has received consulting fees or honoraria from Janssen Pharmaceuticals, GSK, AstraZeneca, and Hoffman-La Roche. JTD holds equity in NuMedii Inc, Ayasdi, Inc. and Ontomics, Inc. No writing assistance was utilized in the production of this manuscript.

## Notes

https://bitbucket.org/dudleylab/pvs

### Summary of Updates

Minor text/grammar edits. Improved readability.

