## Supplemental Data for "Pleiotropic Variability Score: A Genome Interpretation Metric to Quantify Phenomic Associations of Genomic Variants" (from another version of this preprint: version 1, 2021-07-19)

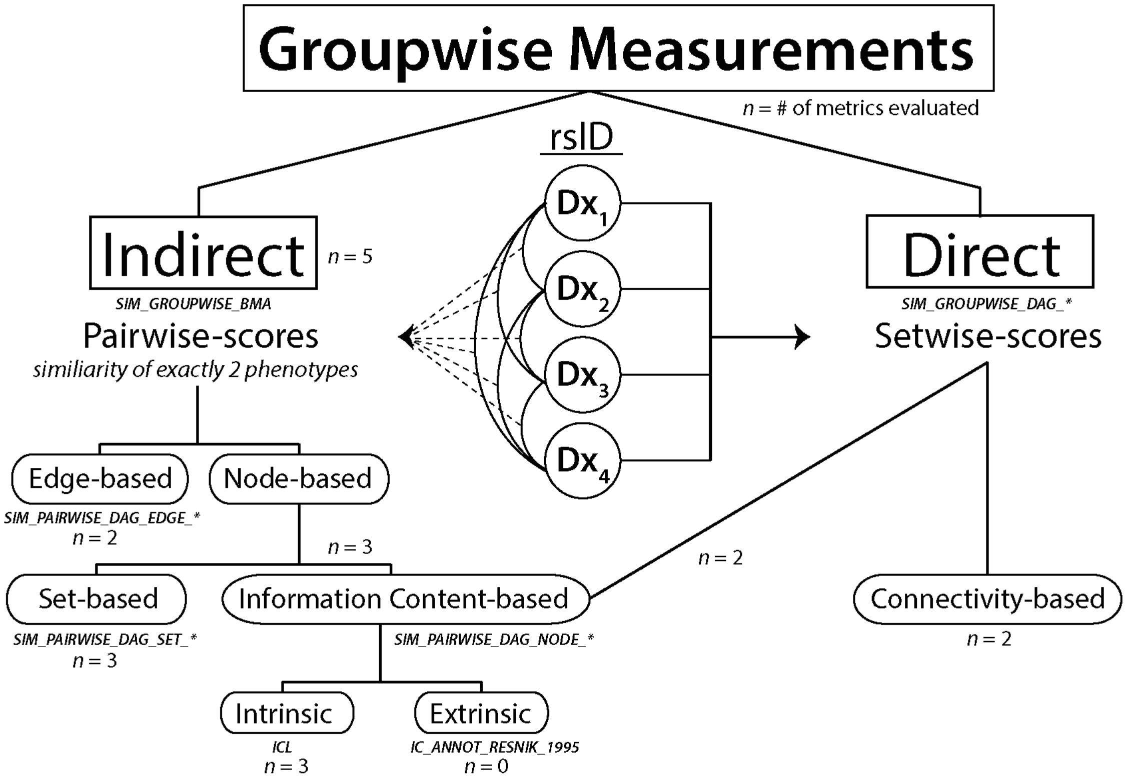


**Figure S1:** Combination of semantic similarity methods assessed to derive final set of scoring methods to compute pleiotropy
Information content based metrics: intrinsic (sanchez, zhou, seco) and extrinsic (resnik); Pairwise metrics: IC-based (resnik,lin, schilcker, jc)
Groupwise metrics: direct (to, nto, ui, lp, lee, ali_and_deane and gic) and Indirect metrics: min, max, avg, avgnorm, bma, bmm, maxnorm;

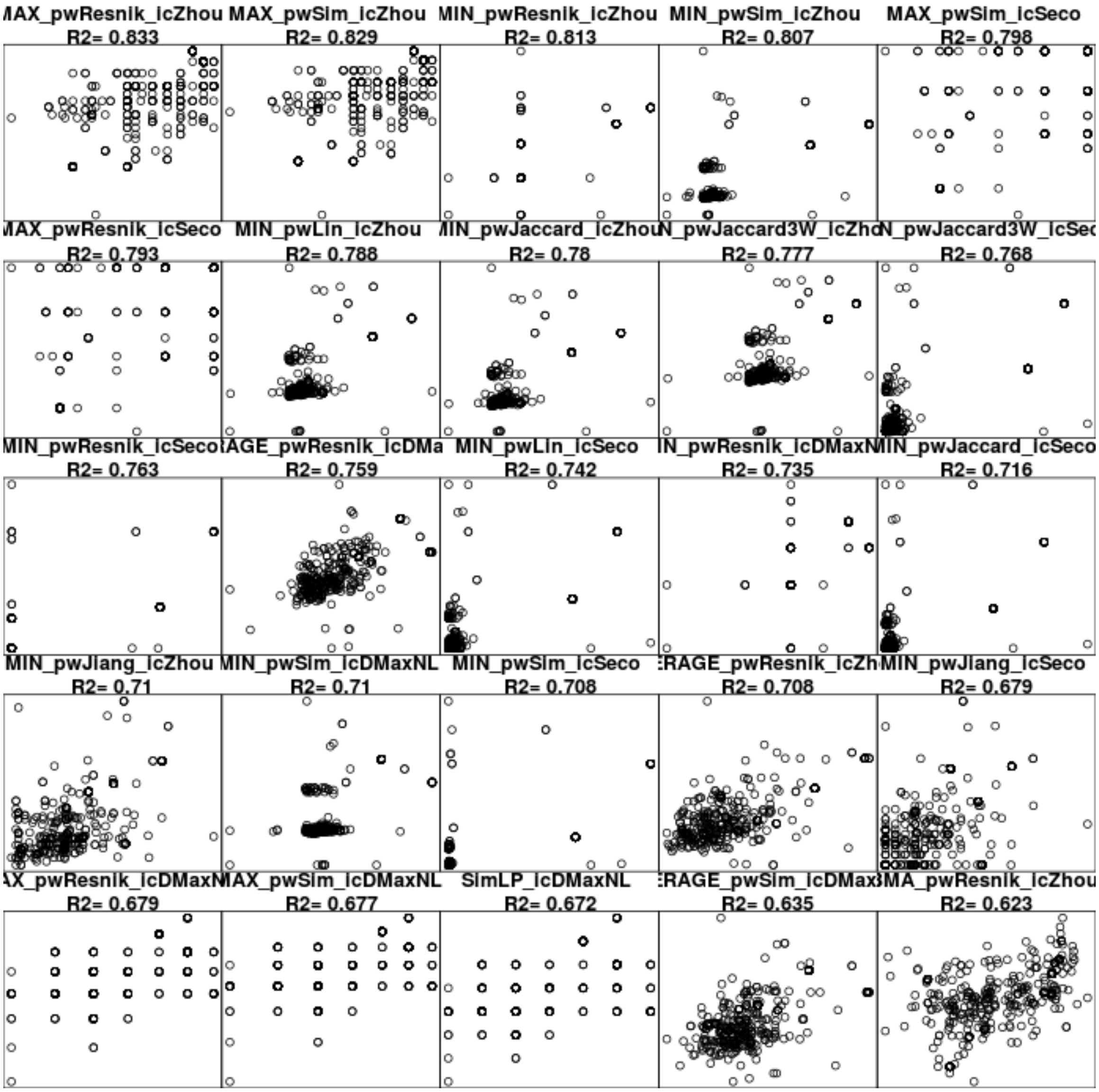


**Figure S2:** Consensus correlation of semantic similarity metrics derived using HPO and DO phenotypes. Individual methods names, approximation method(MAX, MIN or BMA), and indication of grouping the terms (information content, pairwise, group wise or indirect) are provided in above individual plots.

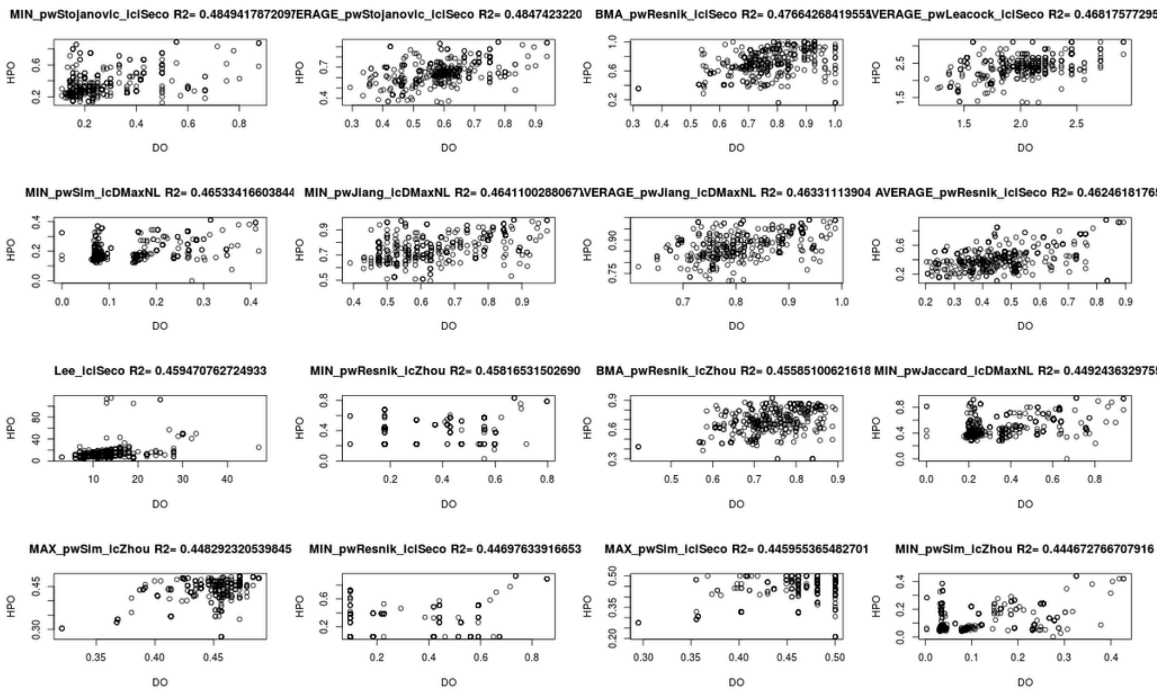


**Figure S3:** Consensus correlation of16 methods using SNPs with more than two phenotypes (phenotype length=2)

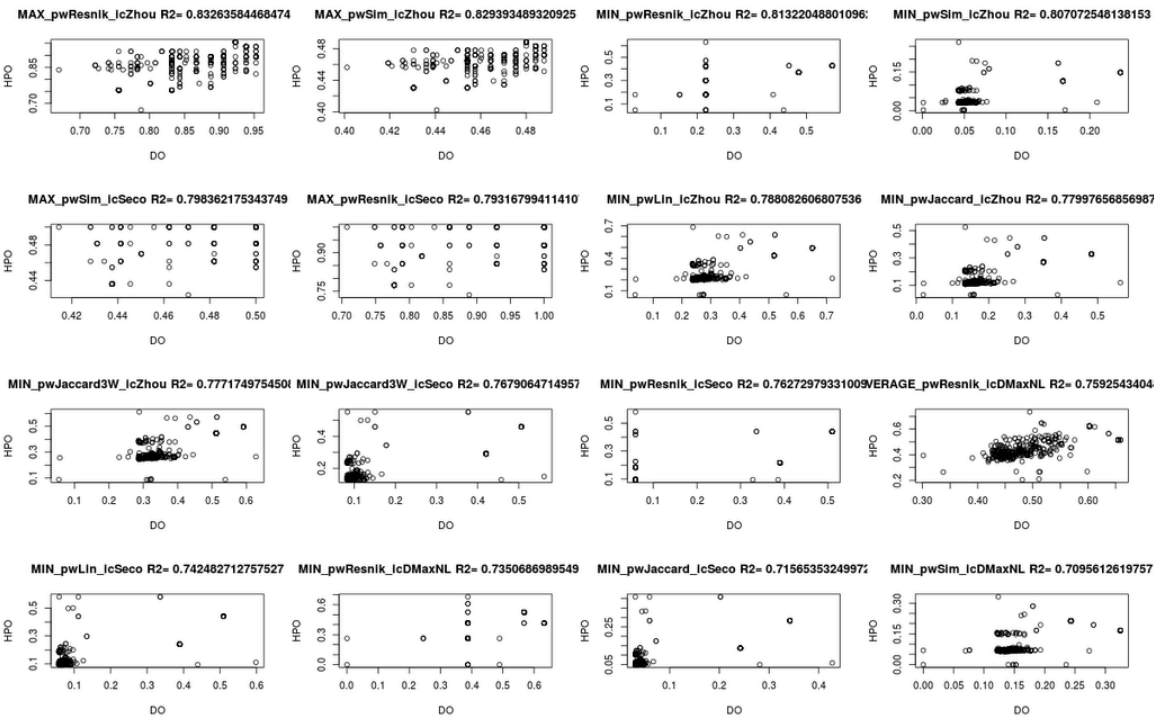


**Figure S4:** Consensus correlation of top 16 methods using SNPs with more than four phenotypes (phenotype length=4)

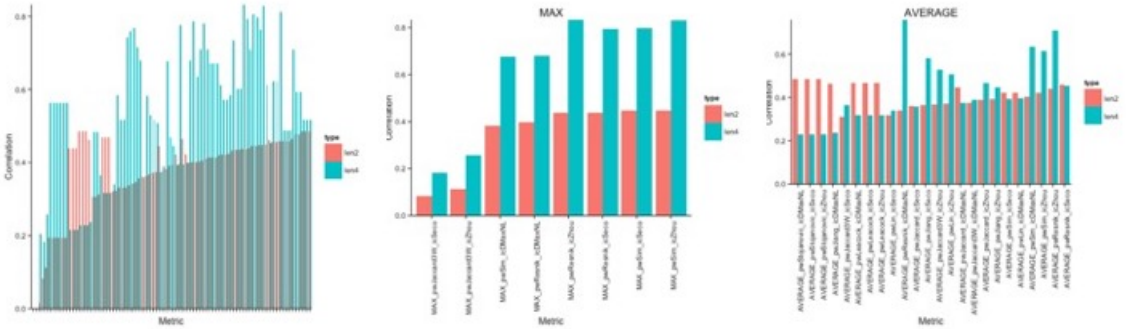


**Figure S5:** Semantic similarity assessment using *Min, Max, Average* approximations using variants with two phenotypes (len2) and four phenotypes (len4)
Data to generate these plots are provided in Supplementary Data: PVS-Phenotype_Len_Correlations.xlsx

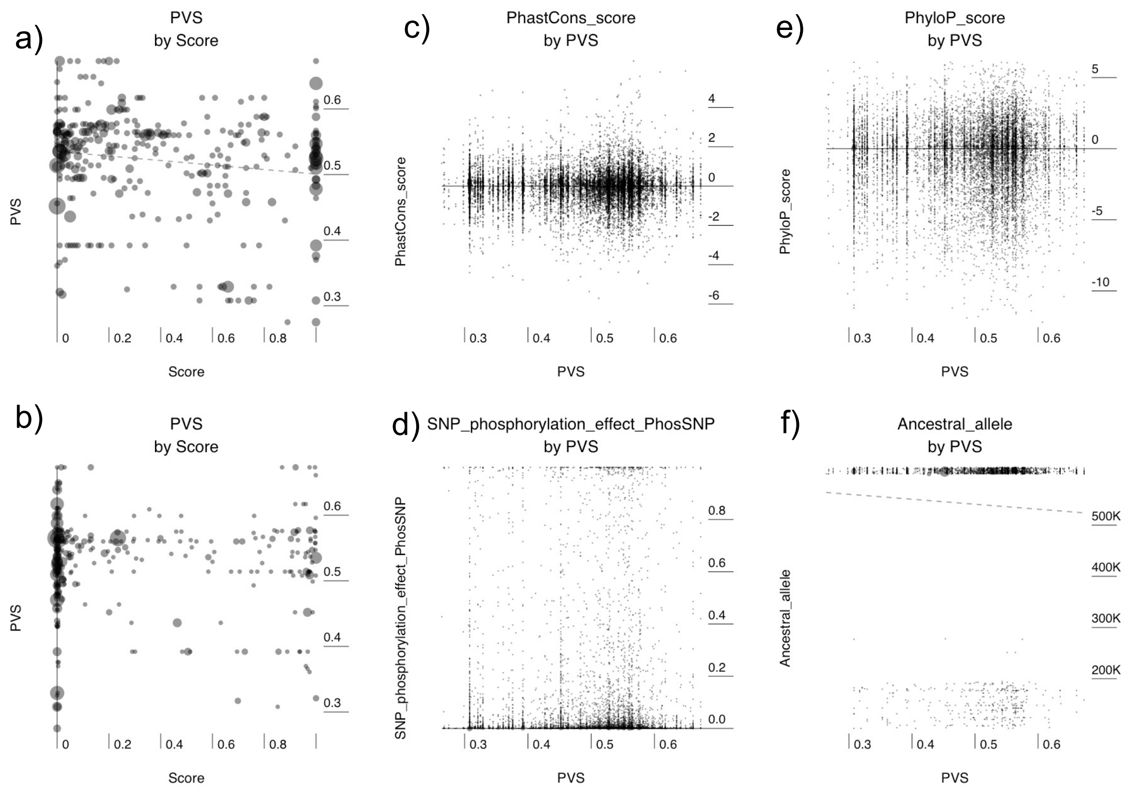


**Figure S6:** Pairwise correlations of PVS with a) SIFT b) PolyPhen 2 c) PhastCons d) PhoSNP e) PhyloP and f) ancestral allele
